# Perturbation of BRMS1 interactome reveals pathways that impact cell migration

**DOI:** 10.1101/2021.02.04.429764

**Authors:** Rosalyn Zimmermann, Mihaela E. Sardiu, Christa A. Manton, Md. Sayem Miah, Charles A.S. Banks, Mark K. Adams, Devin C. Koestler, Douglas R. Hurst, Mick D. Edmonds, Michael P. Washburn, Danny R. Welch

## Abstract

Breast Cancer Metastasis Suppressor 1 (BRMS1) expression is associated with longer patient survival in multiple cancer types. Understanding BRMS1 functionality will provide insights into both mechanism of action and will enhance potential therapeutic development. In this study, we confirmed that the C-terminus of BRMS1 is critical for metastasis suppression and hypothesized that critical protein interactions in this region would explain its function. Phosphorylation status at S237 regulates BRMS1 protein interactions related to a variety of biological processes, phenotypes [cell cycle (e.g., CDKN2A), DNA repair (e.g., BRCA1)], and metastasis [(e.g., TCF2 and POLE2)]. Presence of S237 also directly decreased MDA-MB-231 breast carcinoma migration *in vitro* and metastases *in vivo*. The results add significantly to our understanding of how BRMS1 interactions with Sin3/HDAC complexes regulate metastasis and expand insights into BRMS1’s molecular role, as they demonstrate that BRMS1 C-terminus involvement in distinct direct protein-protein interactions.

## Introduction

Metastasis is a multi-step process that occurs when cells disseminate from the primary neoplasm and eventually colonize distant organs. Successful completion of this complex cascade is associated with nearly all cancer-related morbidities and mortalities. Despite its causative role in cancer-specific mortality and morbidity, a complete understanding of the process and its mechanisms remain elusive. Metastasis is regulated by three types of genes: metastasis-promoting, -suppressing, and -efficiency modifying [1]. The protein of interest in this study, Breast Cancer Metastasis Suppressor 1 (BRMS1), is a metastasis suppressor, which is defined by the ability to suppress metastasis without blocking primary tumor growth [2]. Metastasis suppressors are of distinct interest because, by preventing the successful completion of the metastatic cascade, the devastating sequalae or deaths associated with metastasis would be reduced [3, 4].

BRMS1 was discovered in the year 2000 and its re-expression in multiple cell lines significantly decreases lung metastases in mice [5–8]. In addition, higher expression of BRMS1 in breast, lung, and melanoma cancers is associated with improved patient survival [8–10]. Despite promising functional evidence and clinical correlations, the mechanism by which BRMS1 blocks metastasis is yet to be understood.

BRMS1 is a known member of Sin3 histone deacetylase (HDAC) transcriptional regulatory complexes in multiple eukaryotic cell types [11–13]. It is an established binding partner of SUDS3 and ARID4A and is also associated with several other members of epigenetics-modifying complexes containing SIN3A, SIN3B, HDAC1, and HDAC2 [11]. Relatedly, BRMS1 also exhibits E3 ubiquitin ligase function on the histone acetyl transferase p300, which could also be involved in its regulation of metastatic function [14]. Collectively, it appears that BRMS1 regulation of transcription via epigenetic pathways may be involved in metastasis suppression; however, further studies must be completed to ascribe a definitive mechanism of action.

In order to translate BRMS1 into clinical practice, additional structural and interactome data are necessary. This study leverages previously described features of BRMS1 to clarify its molecular functions within a cell. Previous attempts to define BRMS1 structure-function have been met with uneven success. To date no one has successfully crystallized nor determined structure for full-length, wild-type BRMS1. But computer algorithms and NMR studies have identified structural information for BRMS1 domains [15]. BRMS1 has predicted protein domains that consist of glutamate-rich regions (aa 1-50), two coiled-coil domains (aa 51-81, aa 147-180), and a nuclear localization sequence (aa 198-205) [5]. The N-terminus has been characterized and its 3D structure crystalized [16, 17], but outside of amino acids 51-98 there is no precise protein structure for the entirety of the 246 amino acid BRMS1 protein.

The C-terminus of the BRMS1 protein is of particular interest as it has been shown to play a critical role in metastatic suppression, due to *in vivo* studies demonstrating that alteration of the domain impacts overall metastatic burden [18]. Of note, the serine immediately upstream of the previously characterized critical metastasis suppressor domain was found to be phosphorylated by Cyclin Dependent Kinase 2 (CDK2) [19]. Due to the juxtaposition of S237 to the C-terminal metastasis suppression region, this study focused on identifying proteins with which BRMS1 interacts within that domain as well as testing how phosphorylation within that domain can alter how BRMS1 regulates metastasis suppression, the results of which can be used to better understand how BRMS1 may function to suppress metastasis.

## Results and Discussion

### BRMS1 C-terminal protein structure is important for protein-protein interactions, including Sin3/HDAC complex interaction

Utilizing our knowledge about BRMS1 from previously published functional and interaction studies we generated a panel of BRMS1 mutants: BRMS1^S237D^ (phosphorylation-mimic, hereafter S237D); BRMS1^S237A^ (unable to be phosphorylated, hereafter S237A); BRMS1^1-229^ (lacks critical C-terminal domain); and BRMS1^230-246^ (C-terminal domain with S237 in the center) (**Figure 1A**). Regardless of whether S237 was mutated to aspartic acid or alanine, BRMS1 protein stability was predicted to be compromised using multiple *in silico* prediction methods (**Table 1**). These data suggest that S237 site modifications dramatically destabilize BRMS1 and may be important for overall protein function and should be further investigated.

**Figure 1.**
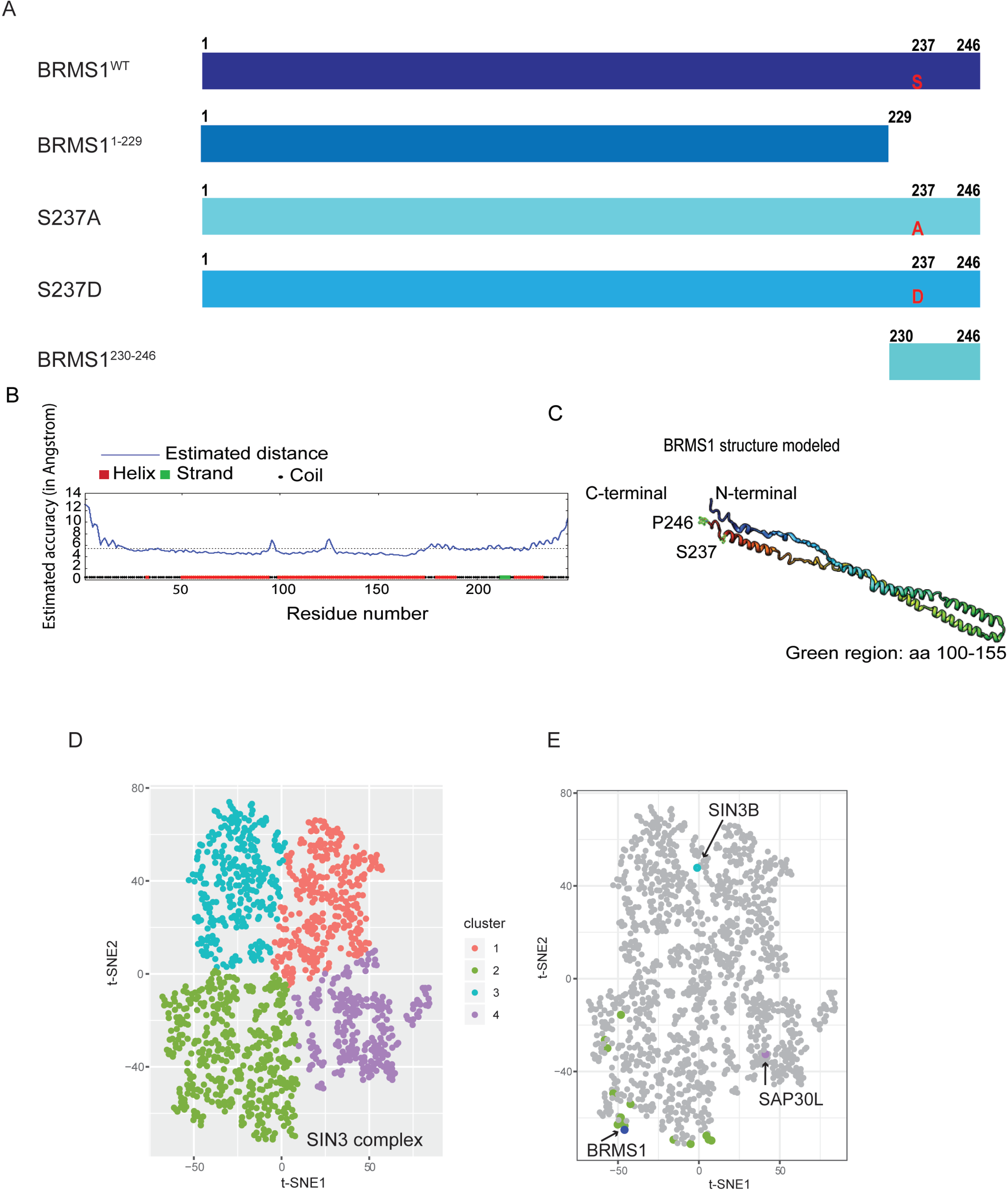
BRM51 C-terminus plays a role in its protein interactome. (A) Mass spectrometry was completed on four BRMS1 mutants with full-length BRMS1 (BRMS1^WT^), BRMS1 aa 1-229 (BRMS1 ^1-229^), BRMS1 (S237A), BRMS1 (S237D), and BRMS1 aa 230-246 (BRMS1^230-246^). (B) Secondary structure prediction of the BRMS1 proteins by I-TASSER, followed by the BRMS1 predicted 3D model (C). The structure generated demonstrates that the C-terminal region consists of helical, coiled, and strand regions. (D) t-SNE analysis was implemented for the analysis of the mutant data. Four clusters are generated by t-SNE. (E). Components of the Sin3/HDAC (in respective cluster colors from Figure 1D) and BRMS1 (in blue) proteins are highlighted.

**Table 1.** Predicted protein stability is impacted by S237 mutations.

| Mutant | Structure | Stability Changes | Predicted Changes | Software |
| --- | --- | --- | --- | --- |
| S237A | Primary | Destabilizing | DDG:<br>-0.214633 | MuProt |
| S237A | Primary | Destabilizing | DDG:<br>-1.69 | I-Mutant |
| S237A | Tertiary | Destabilizing | Folding Rate:<br>-1.1 | Folding Race |
| S237A | Primary | Destabilizing | DDG:<br>-0.095 | DynaMut |
| S237A | Tertiary | Destabilizing | DD Encom:<br>-0.292 kcal/mol | DynaMut |
| S237A | Primary | Order➡<br>Disorder |  | DIM-Pred |
| S237D | Primary | Destabilizing | DDG:<br>-0.0552 | MuProt |
| S237D | Primary | Destabilizing | DDG:<br>-1.0 | I-Mutant |
| S237D | Tertiary | Destabilizing | Folding Rate:<br>-1.8 | Folding Race |
| S237D | Primary | Destabilizing | DDG:<br>-0.457 | DynaMut |
| S237D | Tertiary | Destabilizing | DD Encom:<br>-0.353 | DynaMut |
| S237D | Primary | Order➡<br>Disorder |  | DIM-Pred |

Predictive software (listed) suggests that S237 plays a role in BRMS1 overall structure.

Given the lack of crystal structure for BRMS1 (and specifically the C-terminus), we used the I-Tasser structure prediction method to predict the three-dimensional structure of BRMS1 using its amino acid sequence (**Figure 1B** **and** **1C**). The algorithm-predicted structure identifies that the C-terminal region may consist of helical, coiled, and strand regions, and the phosphorylation site of interest lies within a coiled domain (**Figure 1B**). The predicted coiled domain, coupled with a previous classification of this region as intrinsically disordered - both of which are associated with protein-protein interactions - supported our hypothesis that the region is involved with protein binding and that the interactome is involved in metastasis suppression [15].

The I-TASSER and *in silico* prediction methods, when combined with our previous findings that S237 near the C-terminus can be phosphorylated compelled us to posit that the phosphorylation site could regulate protein binding. To test this hypothesis, mass spectrometry was completed (**Supplementary Table 1**) in 293T cells transduced with Halo-tagged BRMS1 wild-type (BRMS1^WT^), BRMS1^1-229^, S237A, S237D, or BRMS1^230-246^ (**Figure 1A**). Constructs were precipitated via Halo-tags to prevent any contamination with endogenous BRMS1. These constructs were selected because they allowed focus on both phosphorylation as well as putative protein interaction domain(s). This point is key as BRMS1 contains two coiled-coiled domains which are within BRMS1^1-229^ but which are lacking in BRMS1^230-246^, allowing us to identify proteins binding specifically to each region.

Mass spectrometry identified 1175 proteins whose binding was significantly (QSPEC Z-Score ≤-2) altered in at least one mutant compared to BRMS1^WT^. In order to analyze the significant interacting proteins in an unbiased manner, we used the dimension reduction method t-Stochastic Neighbor Embedding (tSNE), applied to Z-statistics obtained from the QSPEC analysis [20]. This analysis resulted in four distinct clusters (**Figure 1D**, **Supplementary Table 2**). Further inspection identified BRMS1 as a member of cluster two, which contains the majority of the Sin3/HDAC members (**Figure 1E**). This finding reinforces BRMS1 membership within Sin3/HDAC complexes while also refining the interactome by lack of association of BRMS1 with SIN3B or SAP30L, which clustered differently (clusters 1 and 4, respectively) (**Figure 1E**).

### BRMS1 interaction with Sin3/HDAC members is more complex than previously anticipated

To further examine the relationships between the Sin3/HDAC complex members and the BRMS1 mutants, Topological Scoring (TopS) was employed (**Figure 2A**). This method is used in determining a likelihood of binding with each of the proteins across the mutants. The resultant scoring suggests the presence or absence of S237 can play a role in BRMS1 interaction with protein complex members. Intriguingly, though there are some member interactions shared between S237A and S237D (BRMS1, BBX, and HDAC2), overall, the mutants’ likely interaction partners tend to cluster differently (**Figure 2A**). S237A, BRMS1^1-229^, and BRMS1^230-246^ cluster the most similarly. This finding may suggest that in order for BRMS1^WT^ to interact with certain proteins it must include both phosphorylation at S237 as well as additional factors near the N-terminus that BRMS1^230-24^ lacks. That being said, BRMS1^WT^ and S237D have little overlap, sharing only HDAC1, HDAC2, and SIN3A (**Figure 2A**). This could be due to the fact that BRMS1^WT^ could more so exhibit a state of flux in regard to binding partners between its phosphorylated and unphosphorylated state and that it binds to many of these proteins, but this remains merely a hypothesis.

**Figure 2.**
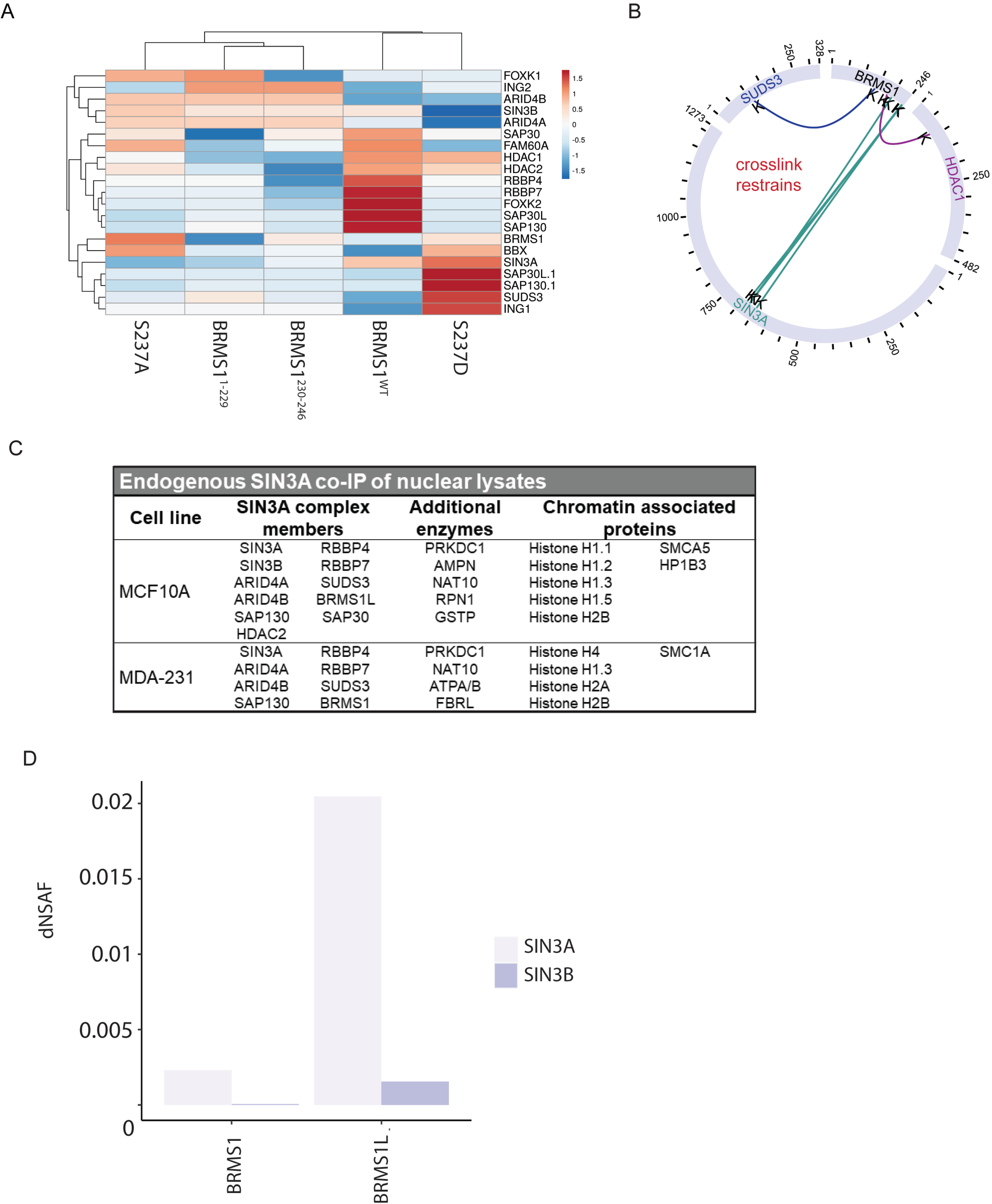
BRMS1 interacts with SIN3/HDAC members alternatively based protein domain. (A) Likelihood of binding for each SIN3A member was evaulation based on TopS score for each of the BRMS1 mutants. Cluster is based upon Eulcidean distance. (B) Crosslink map for the BRMS1 protein, data is extracted from SIN3A XL-MS experiments from Adams et al (23). (C) SIN3A co-immunoprecipation of nuclear lysates followed by MALDI-TOF mass spectrometry. (D) Utilizing MS results from Adams et al (23) dnsAF for SIN3A and SIN3B in relation to BRMS1 and BRMS1 L.

To further examine the Sin3/HDAC complex binding with BRMS1 we utilized recently published crosslinking data of complex members [12]. BRMS1 was confirmed to interact with its known interaction partner SUDS3 (at BRMS1 aa 142), while also binding to the Sin3/HDAC member HDAC1 at BRMS1 aa 184 (**Figure 2B**). BRMS1 also cross-links with SIN3A but not with the related SIN3B (**Figure 2B**). Importantly, the location of SIN3A binding is primarily at BRMS1’s C-terminus (at BRMS1 aa 201, 240, and 242) (**Figure 2B**). These data, in combination with the findings in Figure 1E, suggest that BRMS1 may impact the transcriptional profile of SIN3A complexes greater than SIN3B complexes. In addition, the location of the binding within the C-terminus supports the hypothesis that the C-terminus is involved in determining BRMS1’s protein binding partners. Proximity of the binding sites (aa 240, 242) near S237 further supports the hypothesis that S237 phosphorylation plays a role in regulating BRMS1 binding partners. Taking the data into account with the findings in Figure 2A, the direct cross-link is associated with proteins more greatly associated with S237D (HDAC1,SIN3A, and SUDS3) than the other three mutants (**Figure 2A**) further suggesting the importance of phosphorylation and BRMS1’s interaction within the Sin3/HDAC complex.

To examine the interaction between SIN3A and BRMS1 further, we utilized SIN3A-endogenously expressing cells. SIN3A was co-immunoprecipitated from the nuclear lysates of immortalized, but otherwise normal, MCF-10A breast cell line, as well as within the metastatic MDA-MB-231 breast cancer cell line. These precipitates were then subjected to mass spectrometry through MALDI-TOF analysis. Interactions with SIN3A were found to be context-dependent, i.e. normal breast cell line interactions differ from metastatic cancer cell interactions. SIN3A interacts with several Sin3/HDAC members in MCF-10A cells that it did not interact with in MDA-MB-231 (SIN3B, HDAC2, SAP30, BRMS1L). The only interactor that SIN3A interacted with in MDA-MB-231 that was lacking interaction in MCF-10A was BRMS1. This finding is intriguing as BRMS1L and BRMS1 share high (79% amino acid) sequence similarity [21]. In terms of function, BRMS1L’s role in metastasis is understudied, but two recent studies suggest a role in metastasis suppression [22, 23]. The finding that SIN3A binds to BRMS1 and BRMS1L differently in transformed and non-transformed cells (**Figure 2C**), suggests a role that BRMS1 plays within the Sin3/HDAC complex specifically in cancer that is not apparent in a non-transformed cell line.

To more clearly look at the interactions of BRMS1 and BRMS1L in association with the Sin3/HDAC complexes, we utilized normalized spectral counts of previously published mass spectrometry data [24]. We found a distinct pattern difference of BRMS1 and BRMS1L binding with SIN3A and SIN3B (**Figure 2D**). BRMS1L binds to a much greater extent to SIN3A than BRMS1. Though BRMS1L does not have spectral counts as great with SIN3B as it does to SIN3A, it must be noted that BRMS1 has a very low dNSAF score (5.64 × 10^−5^) with SIN3B (**Figure 2D**). This finding, in combination with the findings in Figures 1E and 2B, compel the hypothesis that BRMS1 interacts with SIN3A over SIN3B. Additionally, the data suggest that BRMS1L may be more relevant in SIN3B complexes, and may also be more common in SIN3A directed complexes than BRMS1, as Figure 2C demonstrates. Taken in combination, these findings hint at a more specific role for BRMS1 within the Sin3/HDAC complex, and leaves room for potential functions independent of its interaction within the complexes.

### C-terminal mutants of BRMS1 disrupt protein interactions and molecular processes

Though significant that BRMS1 mutants altered binding with Sin3/HDAC members, many proteins that exist outside of those complexes also exhibited altered interactions with the mutants. To understand what biological processes may be disrupted due to altered protein interactions, all interacting proteins identified for each mutant were subjected to Reactome pathway analysis (**Figure 3**). Distinct differences in N-terminal (BRMS1^1-229^) versus C-terminal (BRMS1^230-246^) binding were observed. Many of the significant pathways represented in the analysis were not represented in BRMS1^1-229^, e.g., mRNA processing, rRNA processing, and immune responses (**Figure 3**). Many of these same biological processes are disrupted by the C-terminus mutants (S237A, S237D, BRMS1^230-246^), emphasizing the importance of the C-terminus in overall protein function and regulation of protein interactions. Importantly, many of the biological processes are tied to cancer-associated pathologies (i.e. mRNA splicing, immune response) and further study into the BRMS1 C-terminus role in these pathways could identify the molecular role of BRMS1 in metastasis. To identify binding partners distinctly affected by particular domains of BRMS1, all protein interaction changes for each mutant were considered. To ensure that putative interacting proteins were truly associated with BRMS1, an additional criterion was added, i.e., the interactor must have a high TopS value in BRMS1^WT^, to demonstrate an increased likelihood of binding ratio (**Supplementary Table 3**) [25]. Significant overlaps between the C-terminus-associated mutants were observed. All three BRMS1 C-terminus mutants shared 64 proteins whose binding significantly changed compared to BRMS1^WT^ (**Figure 4A, 4B**). These results were in sharp contrast to BRMS1^1-229^, in which those same 64 proteins shared a more similar binding profile to BRMS1^WT^ (**Figure 4B**). Several of the 64 proteins are known promiscuous binders [i.e. eukaryotic translation elongation factor 1 alpha 1 (EEF1A1), heterogeneous nuclear ribonucleoprotein F (HNRNPF)]. To filter promiscuous binders from unique interactors within this dataset, these 64 proteins were subjected to Contaminant Repository for Affinity Purification (CRAPome) analysis [25, 26]. Unique proteins were colored in red in **Figure 4B**. Several of these proteins are associated with cancer, such as TRMT2A (TRNA Methyltransferase 2 Homolog A) and WWP1 (WW Domain Containing E3 Ubiquitin Protein Ligase 1). TRMT2A is a methyltransferase protein and the expression of this gene varies during the cell cycle, with aberrant expression being a possible biomarker in certain breast cancers [27]. Likewise, WWP1 is involved in breast mucinous carcinoma [28]. Two mutants, S237D and BRMS1^230-246^, overlap with the most proteins, with 105 interactors (**Figure 4A**). These proteins were also subjected to CRAPome analysis, which identified several proteins as unique interactors. Two proteins were associated with metastasis, DNA Polymerase Epsilon Subunit 2 (POLE2) and Transcription Factor 2 (TCF2) [29, 30]. POLE2 is associated with circulating tumor cell clusters which, in turn, have been associated with an increased colonization success [29]. TCF2 plays a role in renal cell carcinoma patient progression [31]. In combination, these findings further support the role the C-terminus plays in protein specific binding, while emphasizing the importance of phosphorylation-status on that interaction, and potentially disease state.

**Figure 3.**
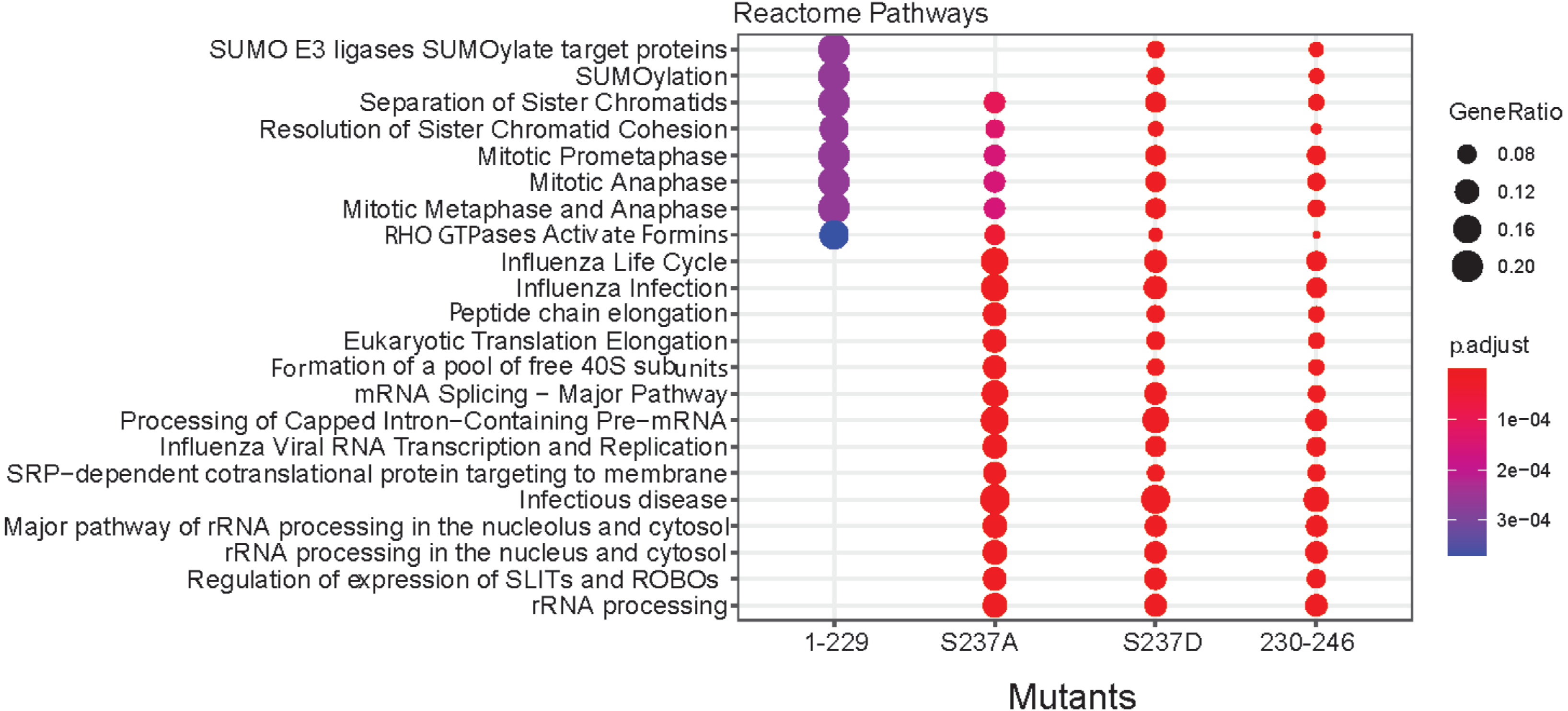
Biological pathway analysis associates BRMS1 mutants with specific molecular processes. The total number of 1175 proteins were subject to the Reactome pathway analysis. Top significant biological pathways with a p-value < 0.01 corresponding to the four mutants are represented in this figure.

**Figure 4.**
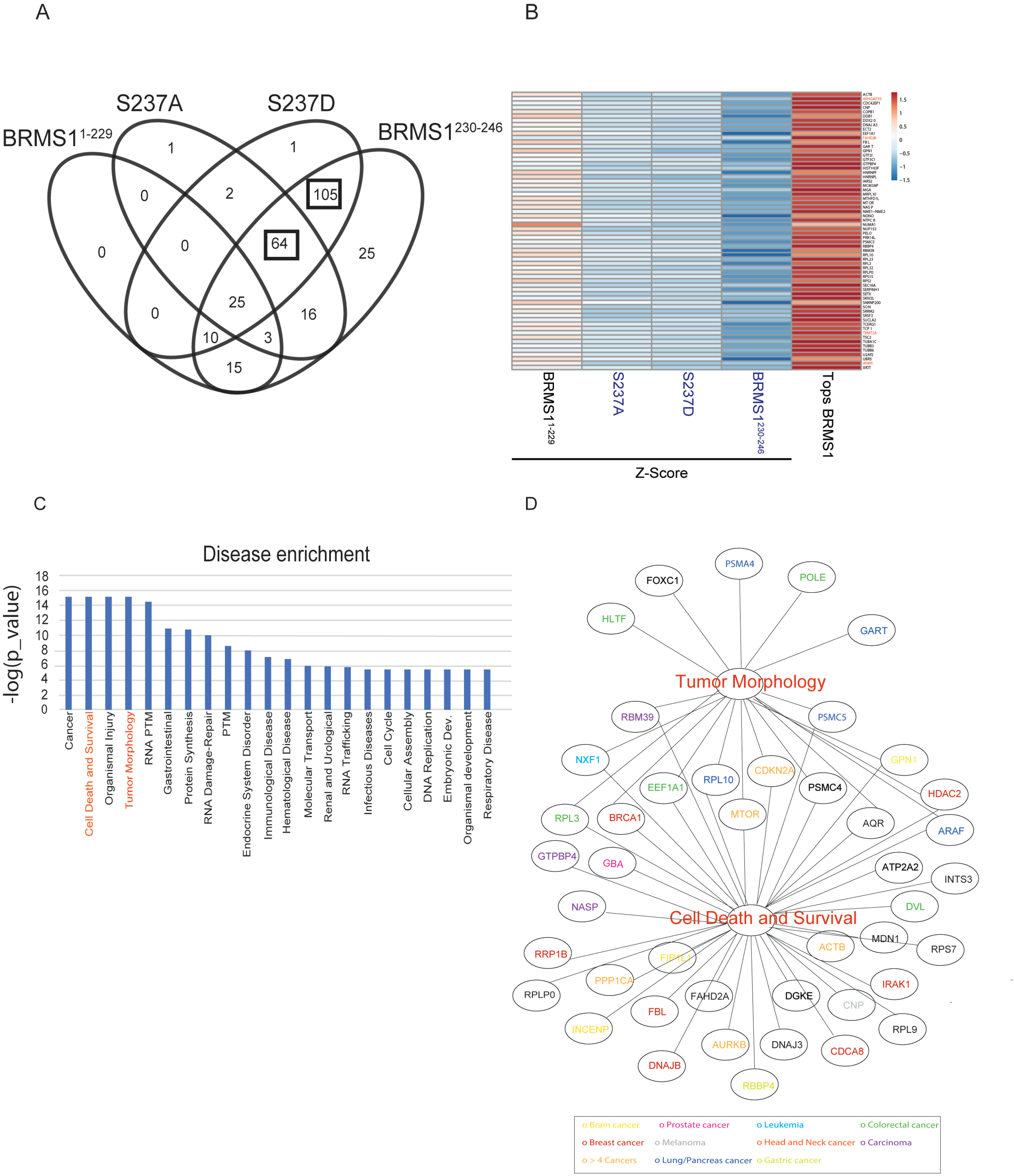
BRMS1 C-terminal binding partners are associated with known pathologies. (A) Venn diagram illustrating the shared proteins between the four mutants. Proteins shared between the three mutants located on the C-terminal and the shared subunits between the S237D and BRMS1^230-246^ are indicated by a black box. (B) The Z-statistics and the TopS values of the 64 proteins shared between the mutants located on the C-terminal are illustrated in (B). The red color corresponds to highest values (TopS or Z-statistics) whereas blue color indicates lower values. Proteins are colored in red if they are in the CRAPome database in less than 3/411 controls with maximum 1 Spectral count. (C) Disease enrichment. Proteins showing significant changes in the mutants in the C-terminal were used in the IPA analysis to determine the most enriched diseases within these proteins. Top 10 enriched classes are displayed in the Figure 4C. (D) Proteins in the Tumor morphology class are illustrated in here. The proteins in the network were color coded according with the type of cancer they associate.

To further characterize how changes in interactor binding may play a role in the development of certain pathologies, disease enrichment within Ingenuity Pathway Analysis (IPA) was completed (**Figure 4C** and **Supplementary Table 4 (Worksheet 2)**). This analysis included all interactor proteins with the C-terminus mutants (S237A, S237D, and BRMS1^230-246^). Several of the enriched pathologies are often associated with neoplasia, including cell cycle dysregulation, cell death and survival, RNA damage repair, embryonic development, immunological disease, and infectious disease. The latter findings compliment previously published results in which re-expression of BRMS1 in metastatic MDA-MB-435 cells were highly enriched for upregulated immune response genes [32].

Two disease-associated categories were enriched and of distinct interest. The first, tumor morphology, is of interest as previous studies showed that BRMS1 re-expression alters expression of the cytoskeletal proteins Focal Adhesion Kinase (FAK), Src, and Fascin [33–35]. These proteins may ultimately change cell morphology and motility [36], which are essential for cancer metastasis [1] (**Figure 4D**). Additionally, enrichment for cell death and survival pathways mirror previous findings as BRMS1 is associated with inducing apoptosis in prostate and non-small cell lung cancers [37, 38]. BRMS1 is also associated with HDAC1 and NF-κB, both of which regulate apoptosis [38].

Within these two enriched disease pathways, several proteins overlap, including BReast CAncer gene 1 (BRCA1), mechanistic target of rapamycin (mTOR), and Cyclin Dependent Kinase Inhibitor 2A (CDKN2A) (**Figure 4D**). Several of the proteins, shared or not, have been associated with many cancers (**Figure 4D**). Proteins that are associated with the C-terminus of BRMS1 strongly suggests that the phosphorylation status of S237 plays roles not only in how BRMS1 functions within the cellular environment, but a distinct role that may shape the metastatic potential. This conclusion is consistent with previous reports that BRMS1 alters multiple steps in the metastatic cascade [33–35, 39].

### BRMS1 domain and phosphorylation status may impact metastatic gene association

To test the impact of phosphorylation site presence or status on known metastasis-associated genes we measured expression changes on genes previously published to be regulated by BRMS1 [35, 40–43] (**Figure 5A**). To better translate findings from Figures 3 and 4 we completed qRT-PCR in 293T cells that were transduced with BRMS1^WT^ and its mutants (**Supplementary Figure 1**). As these measurements were performed in a non-cancerous cell line, many of the changes that have been reported in cancer cells were not expected to be identical. For example, phenotypes necessary for the cells’ metastatic features would be expectedly low so that measurement of BRMS1 diminishment of expression from baseline would be insignificant. This is true for Fascin, PI4K2NA, and RhoC, all of which would be expected to be downregulated, but had no changes within 293T cells (**Figure 5A**). The only significant change was in miR-10b, which was downregulated in both cell lines [43]. Interestingly, miR-10b was only significantly decreased in S237D and BRMS1^230-246^ (**Figure 5A**), suggesting that BRMS1 regulation could be dependent on the presence of S237 and its phosphorylation. This result explains previous findings that BRMS1 C-terminus is responsible for regulating miR-10b expression and points to BRMS1 phosphorylation as a key factor that does so [18]. Further studies must be done to understand the importance of BRMS1 regulation of miR-10b, but the finding does suggest a potential role for S237’s regulation of metastasis.

**Figure 5.**
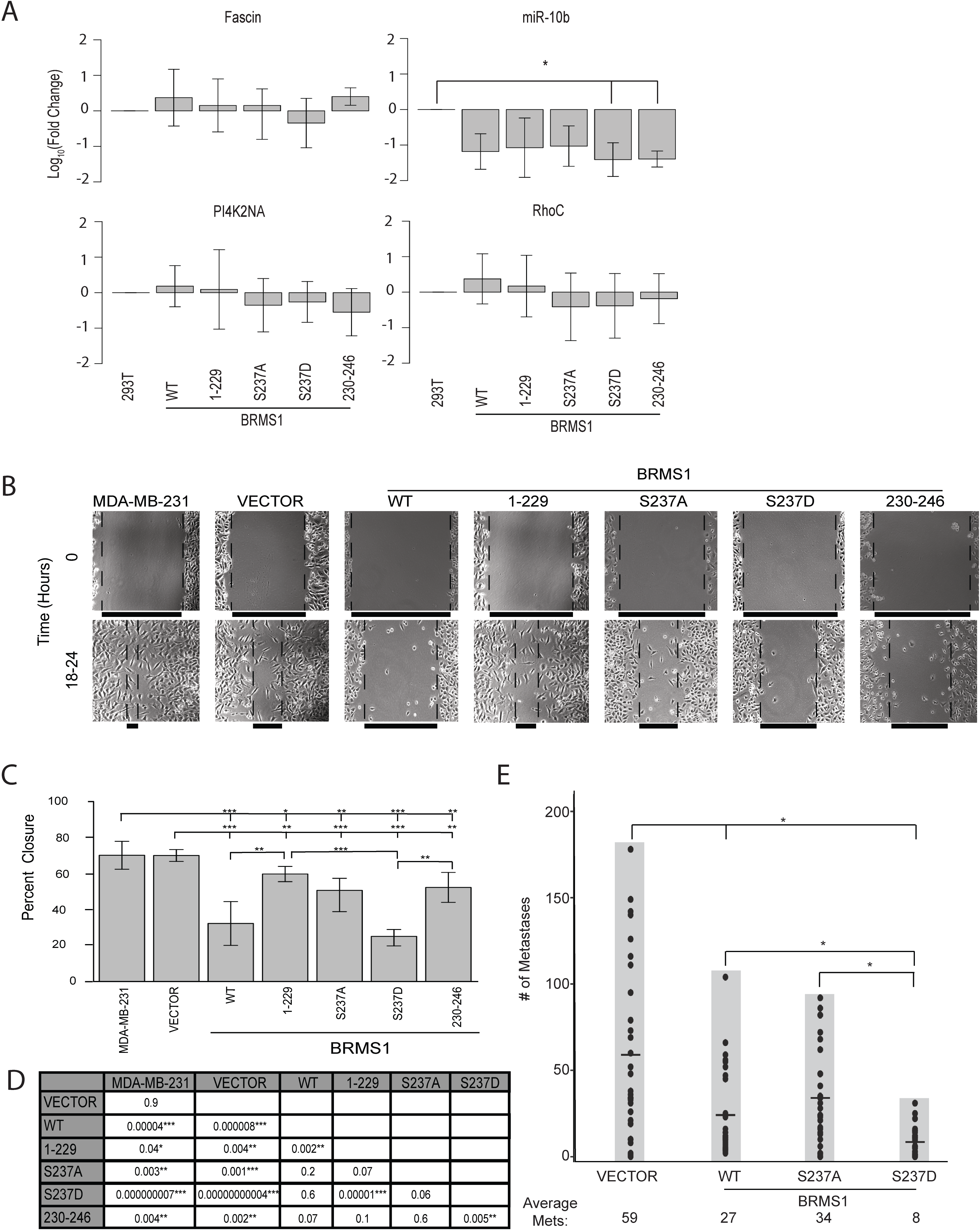
BRMS1 Phosphorylation is associated with a metastasis suppression phenotype. (A) Average qRT-PCR results for 293T cells transduced with BRMS1^WT^, BRMS1^1-229^,S237A, S237D, BRMS1^230-246^. Quantification includes 3 clones per construct and an n=3. Samples were analyzed with a Kruskal Wallis test, followed by a Dunn’s adjustment. * indicates p -values ≤ 0.05. (B) Representative images for migration analysis on parental MDA-MB-231, Vector Control transduced MDA-MB-231s, BRMS1^WT^, BRMS1^1-229^, S237A, S237D, and BRMS1^230-246^ MDA-MB-231 transduced cells. (C) Quantification of migration assays, analyzed with a Kruskal Wallis test, followed by a Dunn’s adjustment. *, **, and *** indicate p-values of ≤ 0.05, ≤ 0.01, and ≤ 0.001, respectively. (D) This table includes the p-values for the comparisons completed in (C). (E) IV Injections of MDA-MB-231 cells expressing vector, BRMS1^WT^, S237A, or S237D. Each experimental group represents an n=30. * indicates a p-value ≤ 0.05. Samples were analyzed with a Kruskal Wallis test, followed by a Dunn’s adjustment.

### Domain and phosphorylation status alter MDA-MB-231 migration *in vitro* and lung metastases *in vivo*

To begin assessing whether BRMS1 mutants exert different biological changes in cancer cells, BRMS1 mutants were transduced into metastatic MDA-MB-231 breast carcinoma cells (**Supplementary Figure 1**). Scratch assays were performed to determine the impact of BRMS1 mutants on *in vitro* cell migration. All BRMS1 mutants inhibited migration (**Figures 5B-5D**). Curiously, BRMS1^230-246^ reduced migration slightly more than BRMS1^1-229^, but this comparison was not statistically significant. BRMS1^WT^ and S237D expressing cells were significantly less migratory than BRMS1^1-229^, while only S237D was less migratory than BRMS1^230-246^. These data suggest that the capacity to exist within a phosphorylated state could decrease the migratory capacity. This observation is speculative, as neither S237A nor S237D have a statistically significantly different migratory capacity to BRMS1^WT^, but the wound closure patterns are in opposite directions. Based upon the *in vitro* data, and the lack of significant difference between S237A and S237D, the full impact of phosphorylation-status is unclear but is suggestive of a role it may play in metastasis.

Based upon the migration data, and the definite differences implicating phosphorylation-status, we wanted to determine preliminarily whether BRMS1 phosphorylation was key to BRMS1-regulated metastasis suppression. To test this, we completed intravenous tail-vein injections with MDA-MB-231 cells expressing BRMS1^WT^, S237A, or S237D (**Figure 5E**). Lung colonization for both BRMS1^WT^ and S237D were significantly lower than vector control, while S237A was not. In addition, S237D had a significant decrease in the number of lung metastases than either BRMS1^WT^ or S237A. These findings suggest firstly the inability of BRMS1 to phosphorylate at S237 does impact metastasis suppressor function. Secondly, the degree of significance by which S237D is able to suppress metastasis, with a decrease compared to BRMS1^WT^ and S237A strongly suggests the key role that S237 phosphorylation plays in BRMS1 metastasis suppression.

### BRMS1 and SIN3 members may impact patient survival

As metastasis is the key factor leading to decreased overall patient survival, we questioned if the interactions within the Sin3/HDAC complex could play a role in impacting patient survival. Utilizing data from The Cancer Genome Atlas (TCGA), we examined how BRMS1 interacting partners (**Figure 2B**) as well as its counterpart, BRMS1L (**Figure 2C**), expression correlated in breast cancer patient survival.

Firstly, we compared the expression of the Sin3/HDAC members from normal to breast tumor samples. BRMS1, HDAC1 and SUDS3 significantly increase expression within the tumor samples, while BRMS1L has significantly decreased expression (**Figure 6A**). SIN3A is unchanged. This finding is intriguing as BRMS1 directly binds both HDAC1 and SUDS3, and hints at a potential association with their expression and patient survival.

**Figure 6.**
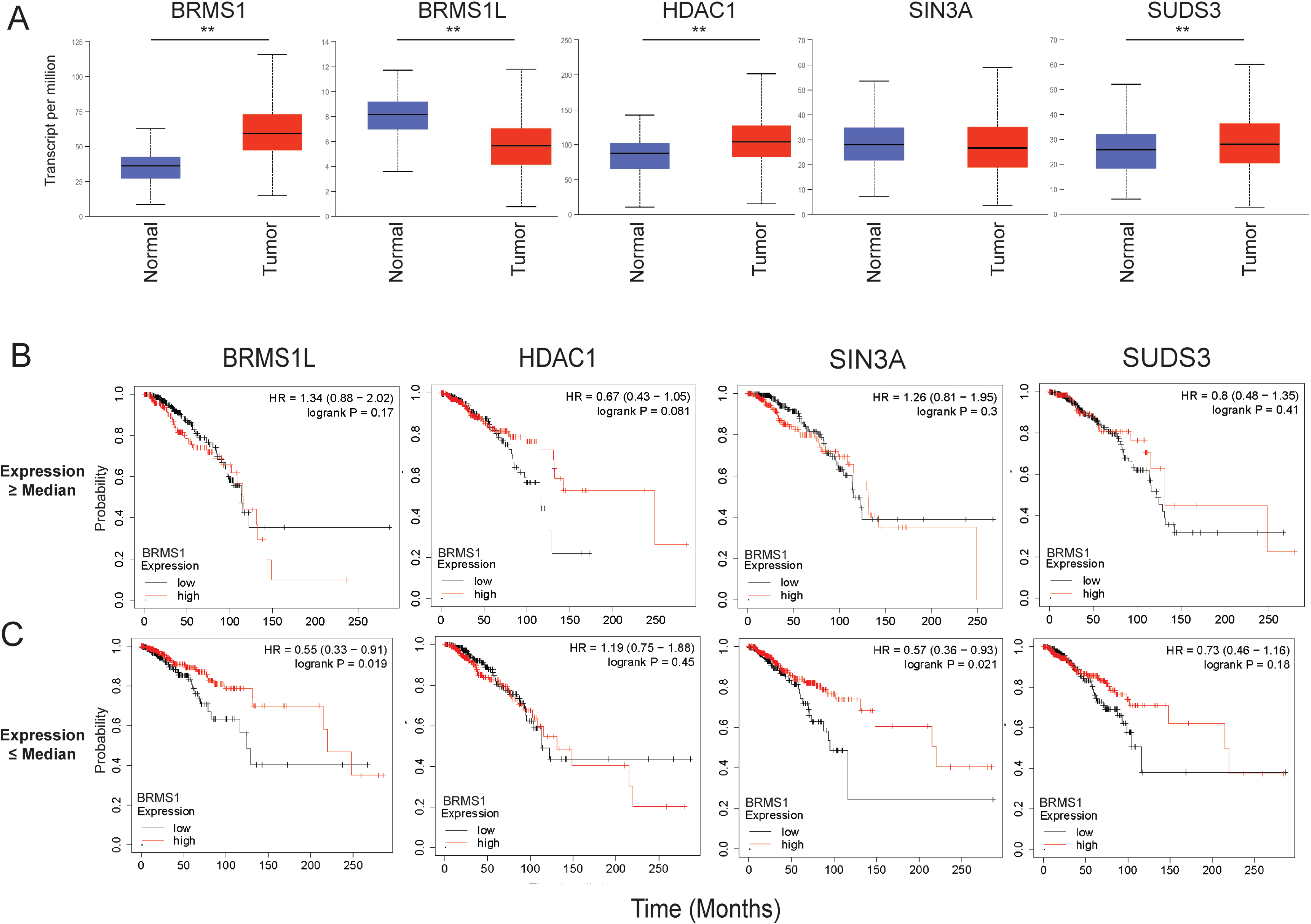
SIN3/HDAC members have altered impact on breast cancer outcomes. (A) Breast Cancer (BRCA) data from TCGA was mined utilzing UALCAN to determine differences in expression of normal compared to tumor tissue.** indicates a p-value < 0.01. (B-C). BRCA data was mined in which SIN3/HDAC members were separated by median expression into those greater than the median (B) or those less than the median (C). BRMS1 expression was then examined within this patients for overall survival, with High BRMS1 (indicated by Red) and Low BRMS1 (indicated by black) were accounted for. This was completed in KM Plotter.

To address this speculation, we also looked for an association between one of the Sin3/HDAC members and BRMS1 expression. To do this we first separated patients by expression of each Sin3/HDAC BRMS1-interacting member by expression greater than or less than the median expression of that particular member. Within those subsets BRMS1, expression (high vs low) was examined for patient overall survival (**Figures 6B, 6C**). When looking across all Sin3/HDAC members associated with a high median expression cohort, BRMS1 does not appear to play a role in increased patient survival (**Figure 6B**). In fact, we only see an association with patient survival in the low expression cohorts of BRMS1L and SIN3A, in which high expression of BRMS1 is associated with increased patient survival (p-value of 0.019 and 0.021, respectively) (**Figure 6C**). Interestingly, this pattern is similar in the rest of the Sin3/HDAC members, in which across all members in the high-expressing cohort, BRMS1 expression did not play a role (**Supplementary Figure 2**). However, in the low-expressing cohort, BRMS1 high expression plays a role in overall survival for BBX and SAP130 (**Supplementary Figure 3**). As previously noted, BRMS1L appears to play a role in metastasis suppression. It has also been shown the low expression of SIN3A promotes invasion and metastasis, and increased SIN3A expression is associated with increased overall survival [44]. Additionally, both SAP130 (Sin3A Associated Protein 130) and BBX have been directly associated with binding SIN3A [45, 46]. Decreased SAP130 expression has been associated in metastasis previously [47]. These previously published findings, in combination with our results, may suggest that in either low BBX, BRMS1L, SAP130, or SIN3A expressing patients BRMS1 expression will take on the role of a metastasis suppressor, resulting in increased overall survival. However, if BBX, BRMS1L, SAP130, or SIN3A expression is high, BRMS1 expression is not as vital in terms of overall survival as the breast cancer itself should not be highly invasive or metastatic (**Figure 6B**, **Supplementary Figure 2**). Though this data needs further testing to be conclusive it furthers our understanding of BRMS1’s role in relation to the SIN3/HDAC complex, and how this interaction may be key to understanding how this is associated with metastasis.

## Conclusions

This study investigates BRMS1’s C-terminus role in the protein’s function, with an emphasis on the function of the S237 phosphorylation site. We hypothesized, based upon *in silico* prediction methods, that the C-terminus functions in protein binding, and demonstrated through mass spectrometry that the C-terminus can impact BRMS1 associations with Sin3/HDAC complexes as well as proteins outside of the complexes. We model this hypothesis in **Figure 7**, in which we believe that BRMS1 does regulate its anti-metastatic function through altered interactions with proteins within or outside of the complex. This study probed both hypotheses by furthering the understanding of BRMS1 association with Sin3/HDAC by defining direct interactions with SIN3A, HDAC1, and SUDS3. We further define BRMS1’s relationship with SIN3A and determined that it may play a more significant role in SIN3A associated complexes compared to that of SIN3B. We also further defined its relationship with BRMS1L and noticed that, despite sequence similarity, BRMS1 and BRMS1L did not appear to have a mirror image relationship to SIN3A or SIN3B. We also began to probe into the relationship that members of the Sin3/HDAC complex have with BRMS1 that is associated with overall survival.

**Figure 7.**
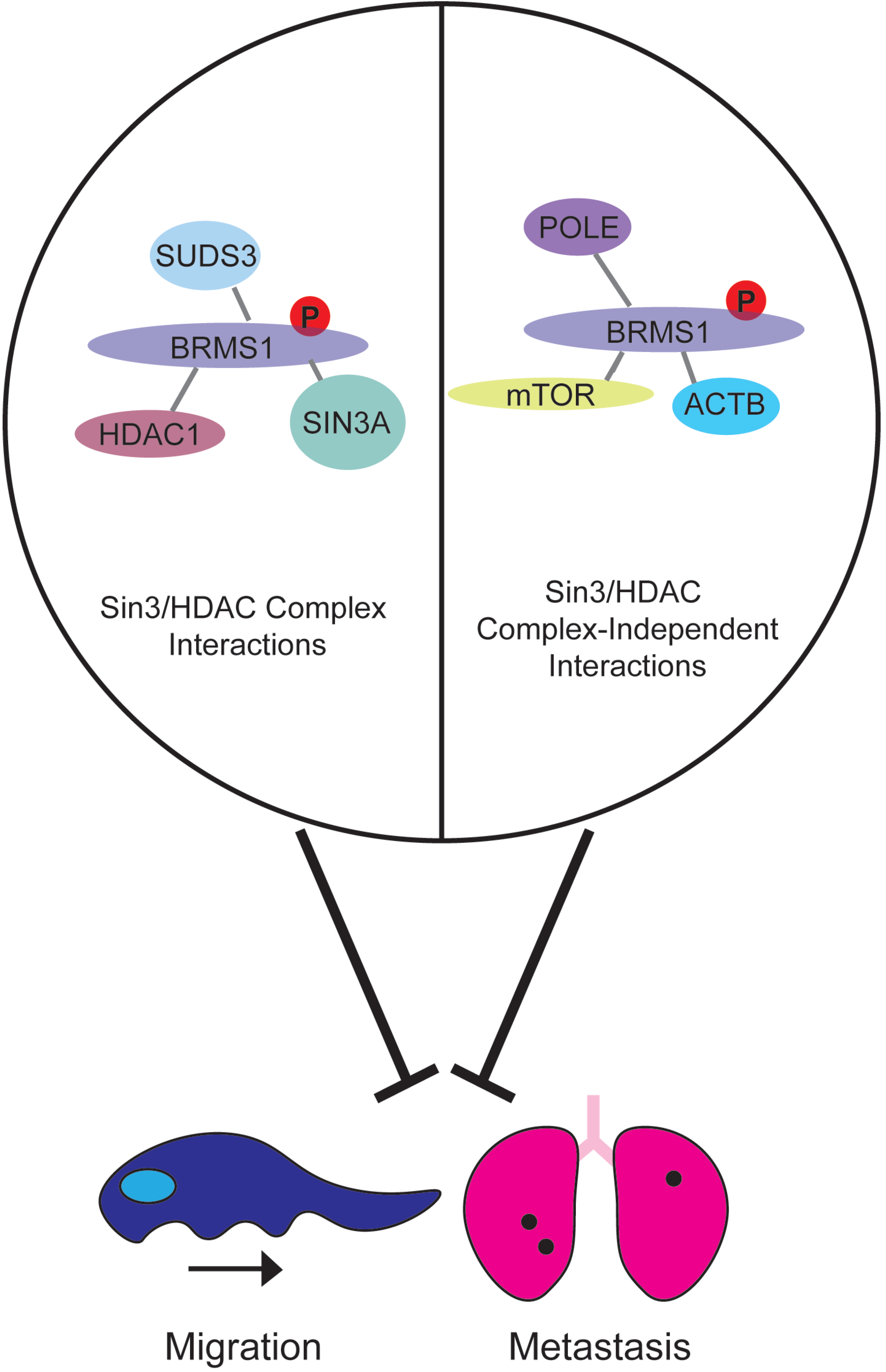
Phosphorylation at S237 contributes to the anti-metastatic phenotype of BRMS1 and alters BRMS1 protein interactions. BRMS1 associates with Sin3/HDAC complex members as well as other proteins which are not part of the chromatin remodeling complexes. We hypothesize that BRMS1 phenotypes are determined by the proteins with which it interacts, but cannot yet ascribe a specific cause-effect relationship responsible for metastasis or migratory suppression.

It is interesting to speculate that once expression of either BRMS1L or SIN3A is lost, it appears that BRMS1 will then function to suppress metastasis and increase survival. Our data begin to define these interactions, in particular with SIN3A, but further studies are required in order to directly define those relationships, including whether the interactions are based on phosphorylation-status to regulate metastasis. The C-terminus and phosphorylation status appear to be vital to regulating the metastasis promoting miRNA miR-10b and corresponding reductions in motility and metastasis. The data also identify specific functionalities of protein complexes previously associated with BRMS1 and metastasis.

This combination of results refine the molecular impact of BRMS1 on regulating metastasis and the potential development of therapeutics based upon BRMS1.

## Materials and Methods

### Cloning of N-terminally Halo-tagged versions of BRMS1 in pcDNA5/FRT

The BRMS1 primers listed in **Supplementary Data** were used to amplify a sequence coding for BRMS1 isoform 1 (NP_056214), or for mutant versions of BRMS1 as indicated in the figure legends, using a previously reported BRMS1 construct as a template [24]. A short synthetic duplex DNA oligonucleotide (described in **Supplementary Data**) was used to clone a short fragment of the C terminus of BRMS1 (amino acids 230-246). DNA fragments were digested with SgfI and PmeI and inserted between SgfI and PmeI sites in pcDNA5FRT-Halo [12].

### Affinity purification of BRMS1 for proteomic analysis

HEK293T cells (1 × 10^7^) were cultured into a 15 cm tissue culture plates for 24 hours then Halo-BRMS1 constructs were transfected using Lipofectamine LTX (Thermo Fisher Scientific). After 48 hours cells were harvested and washed twice with ice-cold PBS. Cells were then resuspended in mammalian cell lysis buffer (Promega) (50 mM Tris·HCl (pH 7.5), 150 mM NaCl, 1% Triton® X-100, 0.1% sodium deoxycholate, 0.1 mM benzamidine HCl, 55 μM phenanthroline, 1 mM PMSF, 10 μM bestatin, 5 μM pepstatin A, and 20 μM leupeptin) followed by centrifugation at 21,000 × g for 10 min at 4 °C. To remove insoluble materials, cell extracts were diluted with 700 μl of TBS (50 mM Tris·HCl pH 7.4, 137 mM NaCl, 2.7 mM KCl) and centrifuged at 21,000 × g for 10 min at 4 °C. Next, cell extracts were incubated overnight at 4 °C with magnetic beads (Magne™ HaloTag® slurry). Before elution, magnetic beads were washed four times with wash buffer (50 mM Tris-HCl pH 7.4, 137 mM NaCl, 2.7 mM KCl, and 0.05% Nonidet® P40). Proteins bound to magnetic beads were eluted for 2 hours at room temperature using elution buffer containing 50 mM Tris-HCl pH 8.0, 0.5 mM EDTA, 0.005 mM DTT, and 2 Units AcTEV™ Protease (Thermo Fisher Scientific). The eluate was further purified by passing through a Micro Bio-Spin column (Bio-Rad, Hercules, CA) to remove residual beads prior to proteomic analyses.

### MudPIT analysis for BRMS1 and BRMS1 mutant associated proteins

MudPIT analysis for protein identification was previously reported in detail by Banks *et al.* [12]. Briefly, trichloroacetic acid (TCA) precipitated proteins were proteolytically digested with endoproteinase Lys-C and trypsin digestion, respectively. Digested peptides were injected directly into a linear ion trap (LTQ) mass spectrometer using 10-step MudPIT separation approach, then the yielded spectra were collected and identified. Spectra were analyzed using the ProLuCID and DTASelect algorithms. Contrast and NSAF7 software were used, respectively, to rank putative affinity purified proteins according to their distributed normalized spectral abundance values (dNSAF). QSPEC was used to identify enriched proteins in the experimental samples [12].

### Cross-linking Analysis

Cross-linking data were utilized from publicly available data published by Adams et. al 2020 [48]. The BRMS1 cross-links were visualized within the xiView platform [49].

### Topological Scoring

Topological Scoring was completed for all proteins as previously described [25, 50]. Briefly, proteins with significant QSPEC scores were input into the TopS Shiny Application (available at https://github.com/WashburnLab/Topological-score-TopS). This application utilizes the average spectral counts of each bait across all baits to calculate the TopS score, which indicates a likelihood ratio of binding.

### Protein Structure Predictions

BRMS1 protein structure was predicted based upon its amino acid sequence retrieved from NCBI using I-TASSER [51, 52]. Briefly, this program functions predicts, with high accuracy, protein structure based upon primary amino acid sequences by comparing the inputted sequence to known BLAST sequences. This comparison is then used to identify potential protein relatives, followed by threading, in which the LOMETS database utilizes the PDB database to generate alignments. Alignments with sufficient Z-Scores are then excised and assembled. Predicted structures are then subjected to iterative Monte Carlo simulations within a variety of conditions (temperature change simulations, etc.). Predictions with the lowest energy state are selected to be the predictive protein structures before comparing against knowns in the PDB database for structure, ligand binding, and biological function.

### Protein Stability Predictions

The primary sequence of BRMS1 was mutated at S237 to either Aspartic Acid (D) or Alanine (A). These mutants were subject to multiple predictive software applications. MuPro [53] utilizes multiple machine learning techniques to predict the impact a point mutation has on protein stability. I-Mutant [54] is a support vector machine-based tool that functions by utilizing neural networks to predict protein stability changes based upon point mutations. Folding RaCe [55] predictive software utilizes a 790 single point mutant knowledge-based prediction of protein folding rates with a multiple linear regression model approach to predict the impact of single site mutations on folding rates. DynaMut [56] combines normal mode analysis which approximates system dynamics and motions with the mutational analysis, such that the prediction is based upon the impact that the mutation has on overall protein dynamics. DIM-Pred [57] predicts order or disorder of a protein or a protein region, accounting for alterations in amino acid properties, neighboring amino acids residues, and substation matrices.

### Data Analysis

Three biological replicates were performed for each bait protein (i.e. BRMS1 wild-type and four mutants). The distributed normalized spectral abundance factor (dNSAF) [58] was used to quantify the prey proteins in each bait AP-MS. To eliminate potential nonspecific proteins, three negative controls were analyzed from cells expressing the Halo tag alone. A total of 15 purifications were completed and 5085 prey proteins identified (Supplementary File 1). First, wild-type data were compared against the negative control dataset to ensure that non-specific proteins were not included in the analysis (Supplementary Table S1). Second, QSPEC [20] was used on this filtered protein list to calculate Z-statistics between spectral counts measured in wild-type and mutants and determine significant changes in protein levels between these two datasets (Supplementary Table 2). We retained only proteins that had a significant QSPEC Z-statistics of −2 or less in at least one of the mutants. The final group of 1175 proteins that passed these criteria comprised the subunits of the Sin3/HDAC complex and proteins outside the complex.

### t-SNE

To spatially map all significant 1175 proteins, we first applied a T-Distributed Stochastic Neighbor Embedding(t-SNE), a nonlinear visualization of the data followed by a k-means clustering on the two vectors generated from the t-SNE (Supplementary Table 2). The number of clusters used for the k-means was 4. The plot visualization was generated within the R packages stats, cluster, gplots, and ggplot2.

### Co-immunoprecipitation and MALDI-TOF Analysis

To compare the overall composition of SIN3A associated proteins between metastatic and normal breast cells, co-IP of SIN3A (polyclonal Ab generated and validated previously; Lewis et al 2016, Oncotarget, 7:78713, PMCID: PMC5340233) from the nuclear lysates of MDA-MB-231 metastatic breast cancer and MCF10A immortalized but otherwise normal breast epithelial cell lines were analyzed by mass spectrometry. The co-IP samples were electrophoresed followed by in-gel digestion and MALDI-TOF analysis as previously described (Hurst et al 2006; BBRC 348:1429, PMCID: PMC1557677).

### Pathway analysis

In order to determine the biological enrichment of differentially expressed proteins in the mutants, we subjected 1175 proteins to Reactome pathway analysis. Enrichment was completed within the R environment and several packages such as clusterProfiler, ReactomePA, DOSE and enrichplot were used.

### Protein Functional Analysis

Significant proteins were subjected to multiple analyses. Disease enrichment was completed within Ingenuity Pathway Analysis (IPA, http://www.ingenuity.com, Release date: December 2016). Visualization for overlapping significant proteins was completed in Venny 2.1 (https://bioinfogp.cnb.csic.es/tools/venny/). Heatmaps were generated within ClustVis (https://biit.cs.ut.ee/clustvis/).

### Cell Culture

293T cells were cultured in 10% FBS, 0.5 mM NEAA + DMEM/Sodium Bicarbonate (2.438 g/L) cell culture medium (Thermo Fisher Scientific). MDA-MB-231 cells were cultured in 5% FBS, 0.5 mM NEAA 0 + DMEM/Sodium Bicarbonate (2.438 g/L) cell culture medium (Thermo Fisher Scientific). In both 293T and MDA-MB-231s, overexpression cells were first transduced with pENTR followed by pLENTI expression within 293FT cells according to manufacturer’s instructions (Invitrogen). Clones were selected with blasticidin. Each construct contained a single or 3x-Flag epitope tag. Expression was quantified using validated BRMS1 1a5.7 antibody (as described previously [59]), FLAG-M2 antibody (Sigma A8592-IMG), or qPCR for BRMS1^230-246^ using the following primers:

F: 5’- GATCCATGGACTACAAAGACCATGACGGTGATTATAAAGATCATGACATCGATTACAAGGATGACGATGACAAGAAGGCTAGGGCAGCTGTGTCCCCTCAGAAGAGAAAATCGGATGGACCTTGATGATGAC -3’,
R: 5’ - TCGAGTCATCATCAAGGTCCATCCGATTTTCTCTTCTGAGGGGACACAGCTGCCCTAGCCTTCTGTCATCGTCATCCTTGTAATCGATGTCATGATCTTTATAATCACCGTCATGGTCTTTGTAGTCCATG -3’.

Predicted amino acid sequences for each construct are:

BRMS1^WT^
DYKDDDDKMPVQPPSKDTEEMEAEGDSAAEMNGEEEESEEERSGSQTESEEESSEMDDEDYERRRSECVSEMLDLEKQFSELKEKLFRERLSQLRLRLEEVGAERAPEYTEPLGGLQRSLKIRIQVAGIYKGFCLDVIRNKYECELQGAKQHLESEKLLLYDTLQGELQERIQRLEEDRQSLDLSSEWWDDKLHARGSSRSWDSLPPSKRKKAPLVSGPYIVYMLQEIDILEDWTAIKKARAAVSPQKRKSDGP
BRMS1^1-229^
DYKDDDDKMPVQPPSKDTEEMEAEGDSAAEMNGEEEESEEERSGSQTESEEESSEMDDEDYERRRSECVSEMLDLEKQFSELKEKLFRERLSQLRLRLEEVGAERAPEYTEPLGGLQRSLKIRIQVAGIYKGFCLDVIRNKYECELQGAKQHLESEKLLLYDTLQGELQERIQRLEEDRQSLDLSSEWWDDKLHARGSSRSWDSLPPSKRKKAPLVSGPYIVYMLQEIDILEDWTAI
S237A
DYKDDDDKMPVQPPSKDTEEMEAEGDSAAEMNGEEEESEEERSGSQTESEEESSEMDDEDYERRRSECVSEMLDLEKQFSELKEKLFRERLSQLRLRLEEVGAERAPEYTEPLGGLQRSLKIRIQVAGIYKGFCLDVIRNKYECELQ GAKQHLESEKLLLYDTLQGELQERIQRLEEDRQSLDLSSEWWDDKLHARGSSRSWDSLPPSKRKKAPLVSGPYIVYMLQEIDILEDWTAIKKARAAVAPQKRKSDGP
S237D
DYKDDDDKMPVQPPSKDTEEMEAEGDSAAEMNGEEEESEEERSGSQTESEEESSEMDDEDYERRRSECVSEMLDLEKQFSELKEKLFRERLSQLRLRLEEVGAERAPEYTEPLGGLQRSLKIRIQVAGIYKGFCLDVIRNKYECELQGAKQHLESEKLLLYDTLQGELQERIQRLEEDRQSLDLSSEWWDDKLHARGSSRSWDSLPPSKRKKAPLVSGPYIVYMLQEIDILEDWTAIKKARAAVDPQKRKSDGP
BRMS1^230-246^
MDYKDHDGDYKDHDIDYKDDDDKKARAAVSPQKRKSDGP

### Metastasis gene expression analyses with qRT-PCR

293T cells were grown to 90% confluency, collected and RNA extracted an RNA Isolation kit (Zymo Research). Following RNA extraction cDNA completed with the iScript cDNA synthesis kit (Bio Rad). qRT-PCR was completed with a SYBER Green assay, and cDNA loaded at a concentration of 50 ng/uL. The following primers were utilized for analysis:

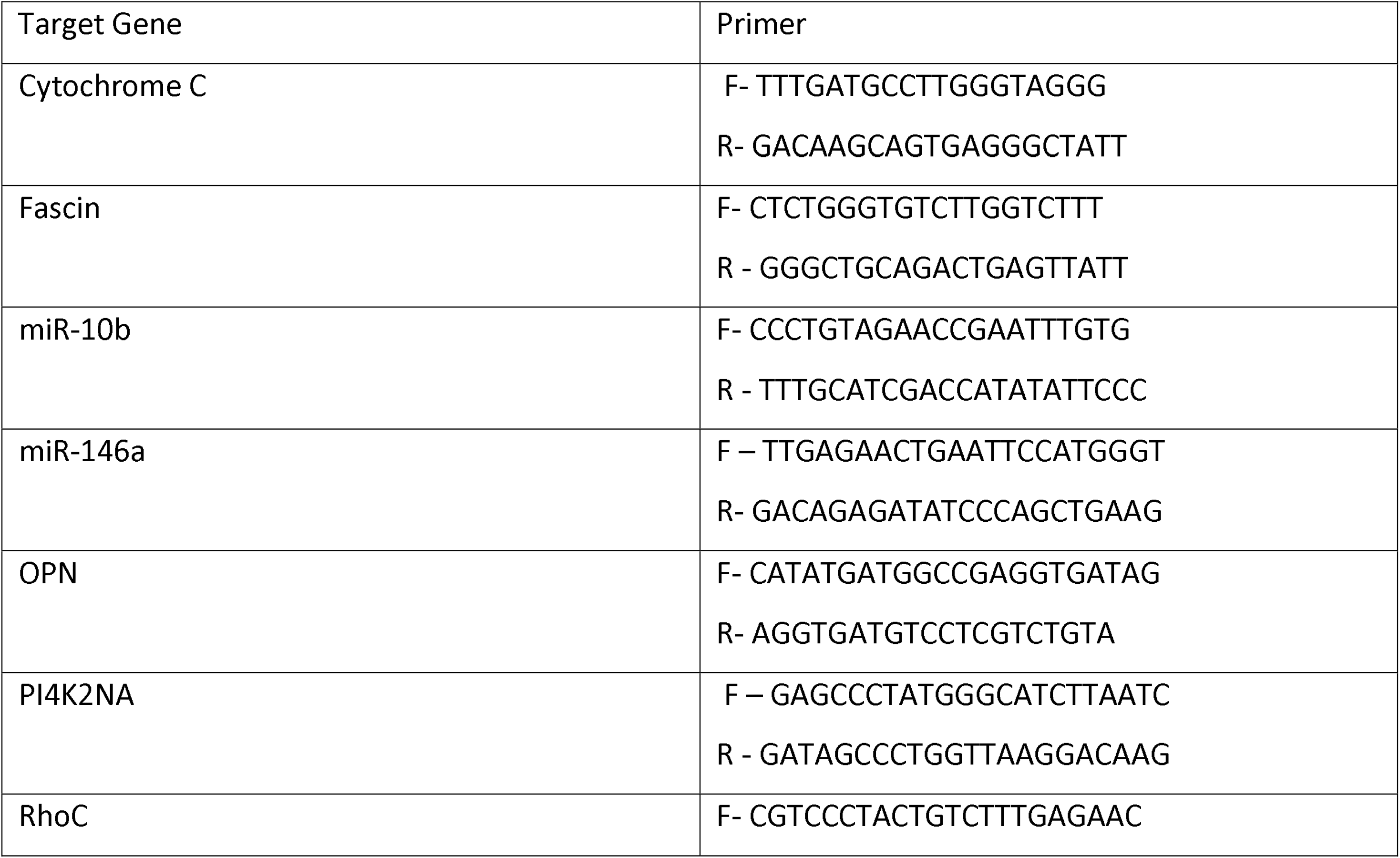
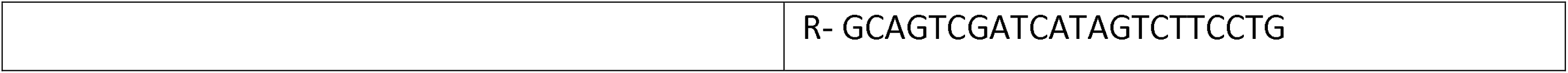

### Migration Assay

Cells were cultured at a seeding density of 750,000 cells/well in 5% FBS, 0.5 mM NEAA + DMEM/Sodium Bicarbonate (2.438 g/L) media in 6-well plates (Thermo Fisher). After 48 hours cells were scratched, washed, and media replaced with serum-free DMEM/Sodium Bicarbonate (2.438 g/L) (Thermo Fisher). Cells were imaged at t = 18 to 24 hrs. Image J (https://imagej.nih.gov/ij/) was utilized for analysis, followed by statistical analysis in R (‘FAS’ package), in which a Kruskal Wallis test followed by a Dunn’s adjustment was completed (https://www.R-project.org/).

### Experimental metastasis assays

To measure lung colonization, three clones of MDA-MB-231 with each BRMS1 construct (BRMS1^WT^, S237A, or S237D) were selected and cells (2×10^5^) were suspended in 100 uL of Hanks Balanced Salt Solution (HBSS, Gibco, #14175-103). Cells were injected intravenously into the lateral tail vein of female athymic mice (Harlan HSD athymic nude-Foxn1nu) aged 3-4 weeks. Lung metastases were allowed to grow for 6 weeks or until the mouse was moribund and required euthanasia. This study was completed using 30 mice and 3 clones per construct. All animal studies were approved by the Institutional Animal Care and Use Committee at the University of Kansas Medical Center (#2014-2208). A Kruskal Wallis test followed by a Dunn’s adjustment was was used for statistical analysis.

### Clinical analyses

The UALCAN web resource was utilized to assess the expression of BRMS1, BRMS1L, HDAC1, SIN3A, and SUDS3 in tumor compared to normal expression (http://ualcan.path.uab.edu/index.html) [60]. All samples were within UALCAN’s BRCA: Breast Invasive Carcinoma dataset. The normal dataset consisted of 114 patients, while the tumor contains 1097. Survival data were obtained from the 2020 version of Kaplan-Meier Plotter breast cancer RNA-sequencing database (https://kmplot.com/analysis/) [61]. Patients were first stratified by the Sin3/HDAC member (i.e. BRMS1L, HDAC1, SIN3A, SUDS3, etc.) by median. Once stratified BRMS1 expression was taken into account, and the p-value is calculated using the log-rank test.

### Visualizations

R packages Gplots, GGplots2, and RColor Brewer were utilized for image analysis of data.

## Funding

This work was supported by the Stowers Institute for Medical Research (MPW, MES, MSM, CAB, MKA); METAvivor Research and Support Inc. (DRW), National Institute for General Medical Sciences [GM112639, F32GM122215 (MPW)]; National Foundation for Cancer Research (DRW), USPHS National Cancer Institute [CA134981, CA168524 (DRW)]; American Cancer Society [PF-16-227-O1-CSM (CAM)]; Susan G. Komen for the Cure [SAC110037 (DRW)]; The K-INBRE Bioinformatics Core supported in part by the National Institute of General Medical Science [P20-GM103418 (RCZ]; the KU Cancer Center Biostatistics and Informatics Shared Resource supported by the National Cancer Institute Cancer Center [P30-CA168524 (DK, DRW)]; and the Kansas Institute for Precision Medicine COBRE supported by the National Institute of General Medical Science [P20-GM130423 (DCK)].

## Data Availability Statement

The mass spectrometry datasets generated for this study are available from the Massive data repository (https://massive.ucsd.edu) using the identifiers listed in Supplementary Table 5. For data generated at the Stowers Institute, links to the original data underlying this manuscript can be accessed from the Stowers Original Data Repository at http://www.stowers.org/research/publications/libpb-1585

## COI

The authors declare no conflicts of interest.

## Author Contributions

RCZ: Conceptualization; data curation; formal analysis; writing-original draft; writing-editing

MES: Formal analysis; writing-original draft; writing-editing

CAM: Conceptualization; funding acquisition; methodology; writing-original draft; writing-editing

MSM: Data curation; formal analysis; methodology; writing-editing

CAB: Data curation; formal analysis; methodology; writing-editing

MKA: Data curation; formal analysis; methodology; writing-editing

DCK: Formal analysis; writing-editing

DRH: Data curation; formal analysis; methodology; writing-editing

MDE: Data curation; formal analysis; methodology; writing-editing

MPW: Conceptualization; supervision; funding acquisition; methodology; project administration; writing-editing

DRW: Conceptualization; supervision; funding acquisition; methodology; project administration; writing-editing

## Notes

https://www.stowers.org/research/publications/libpb-1585

https://massive.ucsd.edu/ProteoSAFe/static/massive.jsp

### Competing Interest Statement

The authors have declared no competing interest.

### Summary of Updates

This manuscript has been updated to reflect our response to reviewers. We are attempting to clarify misinterpreted intent from the previous version of the manuscript, which we believe we addressed in this updated version. In addition, we added more experimental data including Mass Spectrometry, extended interactome data, patient survival analysis, and extended the in vitro and in vivo experiments.

