## Supplemental Figures for "Perturbation of BRMS1 interactome reveals pathways that impact cell migration": Supplemental Figure 1.docx

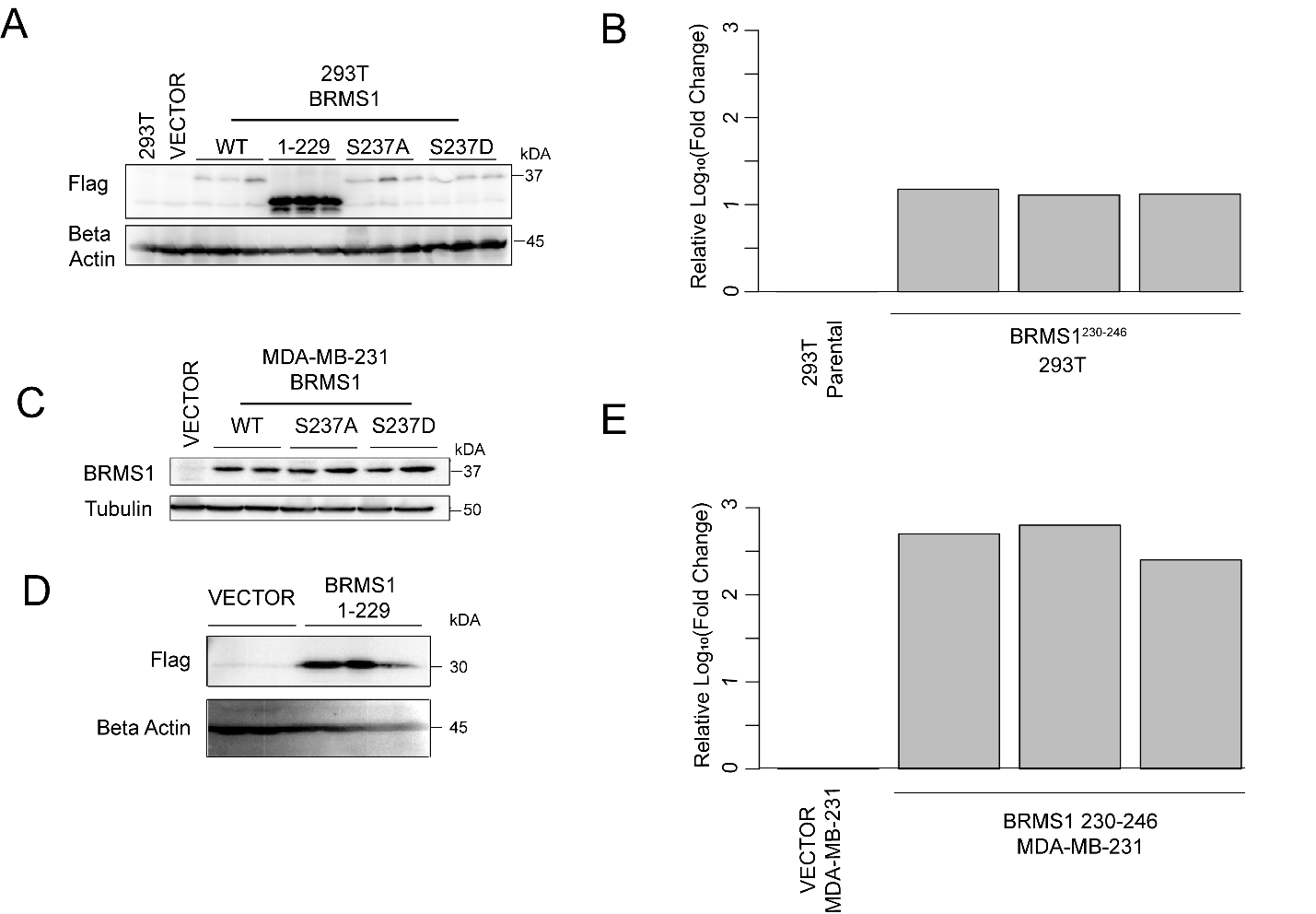

**Supplemental Figure 1 Expression of BRMS1 mutants within MDA-MB-231 cells.** (A) Expression of BRMS1^WT^, BRMS1^1-229^, and phospho-mutants compared to parental and vector control 293T cells. (B) qPCR quantification of BRMS1^230-246^ expression in 293T cells. (C) Expression of BRMS1^WT^ and phospho-mutants compared to vector control MDA-MB-231 cells. (D). Expression of BRMS1^1-229^ within MDA-MB-231 cells. (E) qPCR quantification of BRMS1^230-246^ expression in MDA-MB-231 cells.
