## Supplementary figures and images for "Perturbation of BRMS1 interactome reveals pathways that impact cell migration"

### Suppl Figure 2.tif

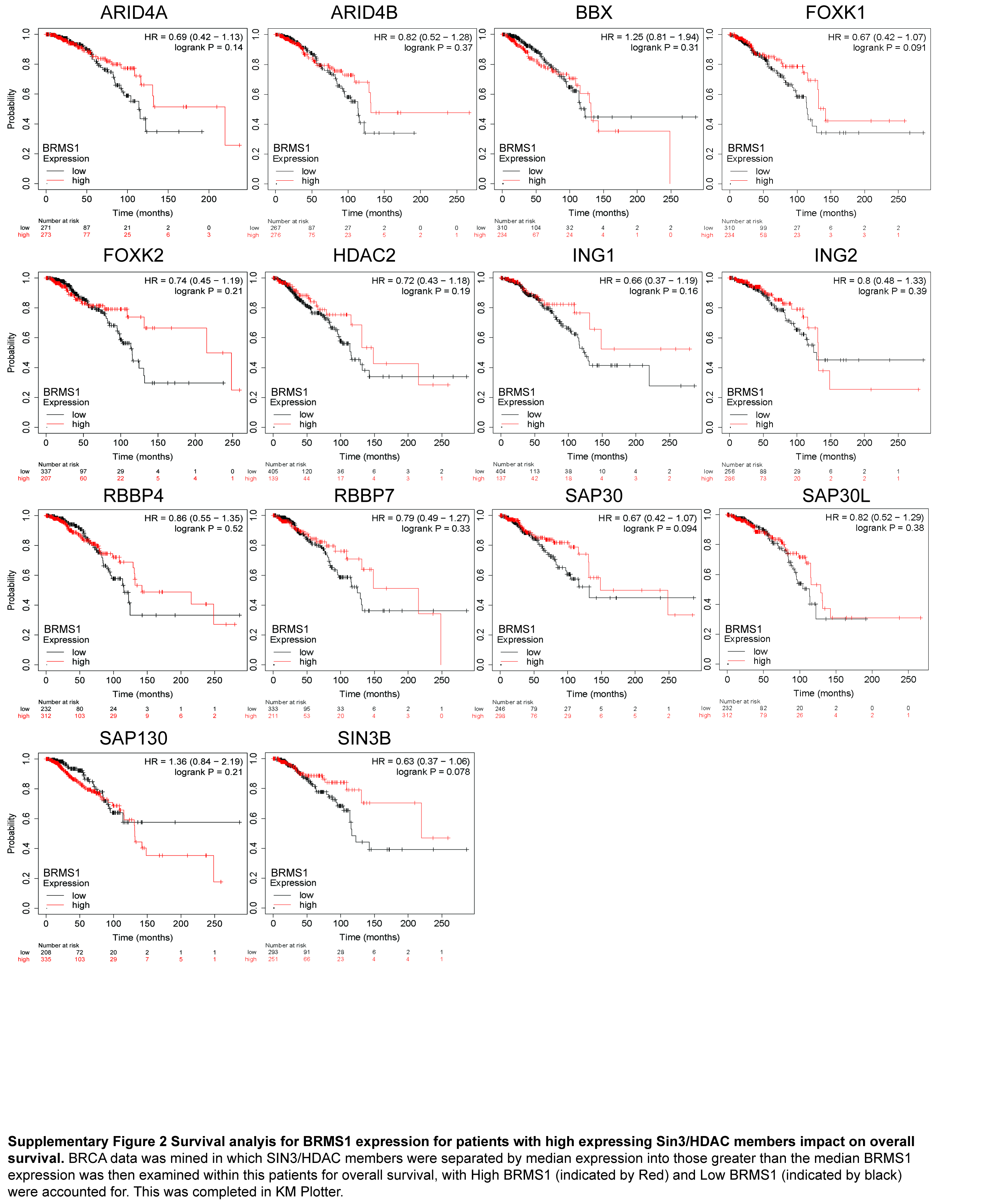

### Suppl Figure 3.tif

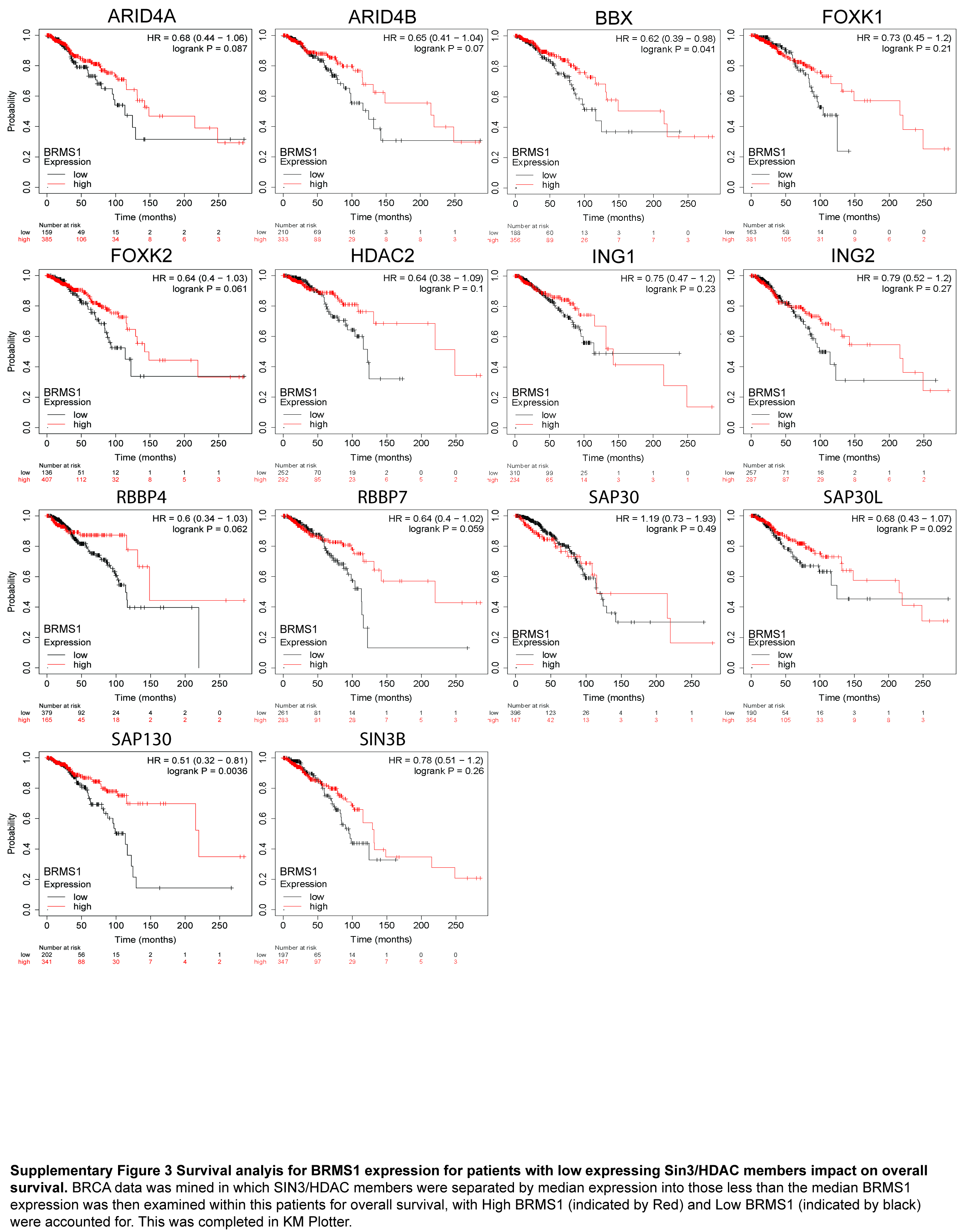
